# Age-related changes in human emotional egocentricity: evidence from multi-level neuroimaging

**DOI:** 10.1101/784215

**Authors:** Federica Riva, Melanie Lenger, Martin Kronbichler, Claus Lamm, Giorgia Silani

**Author notes:** Senior authors who contributed equally to the present work (Claus Lamm, Giorgia Silani). **Corresponding authors:** Claus Lamm, Giorgia Silani.

## Abstract

Emotional egocentric bias (EEB) occurs when, due to a partial failure in self-other distinction, empathy for another’s emotions is influenced by our own emotional state. Recent studies have demonstrated that this bias is higher in children, adolescents and older adults than in young adults. In the latter, overcoming emotional egocentrism has been associated with significant activity in the right supramarginal gyrus (rSMG), as well as increased connectivity between rSGM and somatosensory and visual cortices. Investigations on the neural correlates of EEB in adolescents and older adults are missing. We filled this gap, by asking female participants from three different age groups (adolescents, young adults and older adults, N=92) to perform a well-validated EEB task (Silani et al., 2013) in an MRI scanner. A multi-level analysis approach of MRI data including functional segregation, effective connectivity and structural analyses was adopted. Results revealed higher EEB in older compared to young adults and a comparable EEB in adolescents and young adults. Age-related differences in EEB were associated with differences in task-related rSMG connectivity with somatosensory cortices, especially with S2, which acted as a partial mediator between age and EEB. These findings provide further evidence for the crucial role of the rSMG in self-other distinction in the emotional domain, and suggest that the age-related decline in overcoming EEB is best explained by changes in rSMG connectivity rather than decreased regional activity in that area. This advocates a more systematic investigation of task-related connectivity in studies on aging and life-span development of social-cognitive phenomena.

**Significance Statement:** Empathy comprises both the ability to identify and share another’s emotional state, and the ability to disentangle one’s own from the other’s emotional state. When self- and other-related emotions are conflicting, empathy might be negatively influenced by egocentric tendencies. This phenomenon is referred to as emotional egocentric bias (EEB), with previous research showing that its extent changes across the life-span. Here, we provide evidence that age-related differences in EEB are mainly associated with age-related changes in rSMG effective connectivity, and in particular that higher EEB in older adults is associated to lower rSMG effective connectivity with somatosensory cortices. These findings suggest the importance, particularly in aging, of intact functional connectivity for optimal socio-cognitive functioning.

## 1. Introduction

Emotional egocentric biases (EEB) (Silani et al., 2013) occur when people’s perception of others’ emotions is biased by their own, conflicting emotions – such as, e.g., when someone is not able to fully empathize with a friend’s discomfort for his/her recent break up because he/she has just gotten married. While many different views on empathy exist (Batson, 2009; Singer & Lamm, 2009, for a review), one prevailing definition sees empathy as an isomorphic affective state elicited by seeing or imaging someone else’s affective state, and for which the empathizer is aware that the cause for her affective state lies in the other person’s affective state (e.g., de Vignemont & Singer, 2006). Thus, empathy entails at least two different aspects: i) identifying and sharing the affective state of the other (affective sharing) and ii) keeping separate self-experienced emotions from those experienced by the others (self-other distinction). EEB has been linked to a partial failure in self-other distinction (Silani et al., 2013; Steinbeis et al., 2014; Riva et al., 2016; von Mohr et al., 2019). Whereas some studies have addressed age-related changes in the neural underpinnings of affect sharing (Decety and Michalska, 2010; Chen et al., 2014; Tamm et al., 2017; Riva et al., 2018), less is known about age-related changes in emotional self-other distinction. This gap possibly is due to the paucity of suitable paradigms to measure it. Only recently, we have developed an experimental set-up allowing to detect EEB (Silani et al., 2013), as a proxy of self-other distinction failure. The main idea of this paradigm is to induce either conflicting or matching transient emotional states in pairs of participants, while they are asked to empathize with and to rate the other’s emotions. On the behavioral level, two studies (Steinbeis et al., 2014; Hoffmann et al., 2015) using a similar task approach have shown that children display a higher EEB compared to young adults. Later in the lifespan, by means of a cross-sectional design including four groups (12-17 yrs., 20-30 yrs., 33-55 yrs., 63-78 yrs.) we extended these findings to adolescents and older adults, showing that these two age-groups display a higher EEB compared to young and middle-aged adults. The present study was directly motivated by the observed age-related differences in EEB, with its major aim being to unravel their neural underpinnings.

In this, our main hypotheses were based on previous work performed in healthy young adults (Silani et al., 2013, Steinbeis et al., 2014), which had shown that EEB is predominantly associated with activity in the right supramarginal gyrus (rSMG), and with increased connectivity of rSMG with somatosensory and visual cortices (Silani et al., 2013) and prefrontal areas (Steinbeis et al., 2014). Interestingly, particularly from a lifespan perspective, research on structural brain development showed that specific parts of the parietal lobule (Giedd et al., 1999), notably including the SMG (Gogtay et al., 2004), seem to reach full maturation toward the end of adolescence. Moreover, a linear reduction of gray matter volume of the same area has been observed during adulthood, with a consistent drop after the seventh decade (Courchesne et al., 2000) (Sowell et al., 2003). Taken together, these findings suggest that higher EEB in adolescents and older adults might be linked to age-related changes in structure, function and connectivity of the rSMG.

Therefore, we hypothesized higher EEB in adolescents and older adults to be related to lower recruitment and/or changes in connectivity of rSMG in comparison to a reference group of young adults. Finally, we hypothesized that the predicted differences in brain function are subtended by potential differences in brain structure.

## 2. Materials and Methods

### 2.1 Participants

Ninety-five female participants took part in the study. Subjects were part of three age cohorts: adolescents (AD, 14-17 years; N= 32), young adults (YA, 21-31 years; N=32), and older adults (OA, 56-76 years; N=30). Only female participants were recruited for two reasons: consistency with our previous studies, using the same paradigm, in which we had investigated only females as well, (Silani et al., 2013; Riva et al., 2016), and acknowledged gender differences in empathy and socio-affective skills, including the EEB paradigm we used (e.g.: Schulte-Rüther et al., 2008; Tomova et al., 2014). Differing from our previous work (Riva et al., 2016), we did not add a group of middle-aged adults, since the extent of the EEB in young and middle-aged adults had been indistinguishable in that range. We thus decided to incorporate only younger adults as a reference group against which to compare the adolescent and older-aged group; younger adults instead of middle-aged adults were chosen both because previous imaging work (Silani et al., 2013; Steinbeis et al., 2014) had used this age range as well, and because they are easier to recruit. Three subjects had to be excluded from the analyses (two AD, one YA) for excessive movements (see fMRI analysis paragraph for exclusion criteria) during the scanning session. The final sample thus consisted of 31 AD (age range: 14-17 years old; M= 15.61; SD= 1.03), 31 YA (age range: 21-31 years old; M = 24.52; SD = 2.41) and 30 OA (age range: 56-76 years old; M = 63.42; SD = 4.6). All the participants were right-handed (Oldfield, 1971), had normal or corrected-to-normal vision, and reported no past or present neurological or psychiatric disorder. In addition, the German version of the Mini Mental State Examination (Kessler et al., 2000) was administered to older adults to control for initial stages of neurodegenerative disorders. Only participants with a score higher than 27 (maximum score = 30) were tested, as this cut-off has been reported as most appropriate to screen for cognitive impairment (Kukull et al., 1994). Written consent was provided by participants, who received € 25 each for taking part in the study. The study had received approval by the local ethics committee and was carried out in accordance with the Declaration of Helsinki (latest revision, 2013).

### 2.2 Task and Procedure

The experimental session begun with the participant being introduced to an alleged other participant (a confederate of the study, from now on confederate) and the delivery of the instructions to both of them together, with the aim to get them acquainted. During the instructions, the experimenter explained that the participant and the confederate would have to perform the same task, with the participant lying inside and the confederate seated outside the MR scanner. After this initial phase, the participant was accompanied to the MR scanner room. Overall, the participants had to complete three tasks in the scanner: an empathy task, the EEB task, and an imitation-inhibition task (Brass et al., 2005). The present paper focused on the results of the EEB task only, while the results of the other tasks will be or have been reported elsewhere (Riva et al., 2018).

The EEB paradigm implemented in the current study closely followed the procedure of the second fMRI experiment described in Silani et al. (Silani et al., 2013). Each trial of the EEB task comprised a stimulation phase and a rating phase. In the stimulation phase, transient pleasant or unpleasant affective responses were induced by means of visuo-tactile stimulation of the participants. In one run, they were instructed to empathize with the feelings of the other participant (i.e., the confederate). The affective responses elicited in the pairs of participants could be either congruent (both pleasant or both unpleasant, *congruent condition) or* incongruent (self pleasant and other unpleasant, or vice versa, *incongruent condition*). The visuo-tactile stimulation consisted of the participants seeing on the screen the picture of an object/animal (e.g., a rose, a snail, maggots) accompanied by the text “YOU” and simultaneously, their left palm was stroked by an experimenter using an object whose tactile qualities resembles those of the displayed object or animal. Next to the first picture, the picture of another object/animal accompanied by the text “OTHER” (see Fig.1) was displayed on the screen, indicating the object/animal with which the confederate’s palm was being stroked. The stimulation phase lasted for 3 s. The visual stimuli were presented and seen by means of a back-projection system installed on the scanner site. Afterwards, participants were asked to rate the pleasantness of the confederates’ feelings. The ratings were provided on a visual analogue scale by moving the cursor on the screen using an MR-compatible response box. The selected screen coordinates were converted offline to a scale ranging from −10 (very unpleasant) to +10 (very pleasant). The participants were instructed to respond as quickly and accurately as possible, with a response time limit of 3 s. Offline, the ratings provided in the *incongruent* condition were compared to the ratings provided in the *congruent* condition. In order to exclude that the difference in the ratings between the incongruent and the congruent other-judgment conditions was due to a mere incongruity effect, another run was implemented, during which participants had to rate their own feelings while undergoing incongruent and congruent stimulation as well (self-judgment run). For both the self- and the other-judgment runs there were 20 congruent and 20 incongruent trials (10 pleasant and 10 unpleasant). Self- and other-judgment runs were counterbalanced between participants.

**Fig 1.**
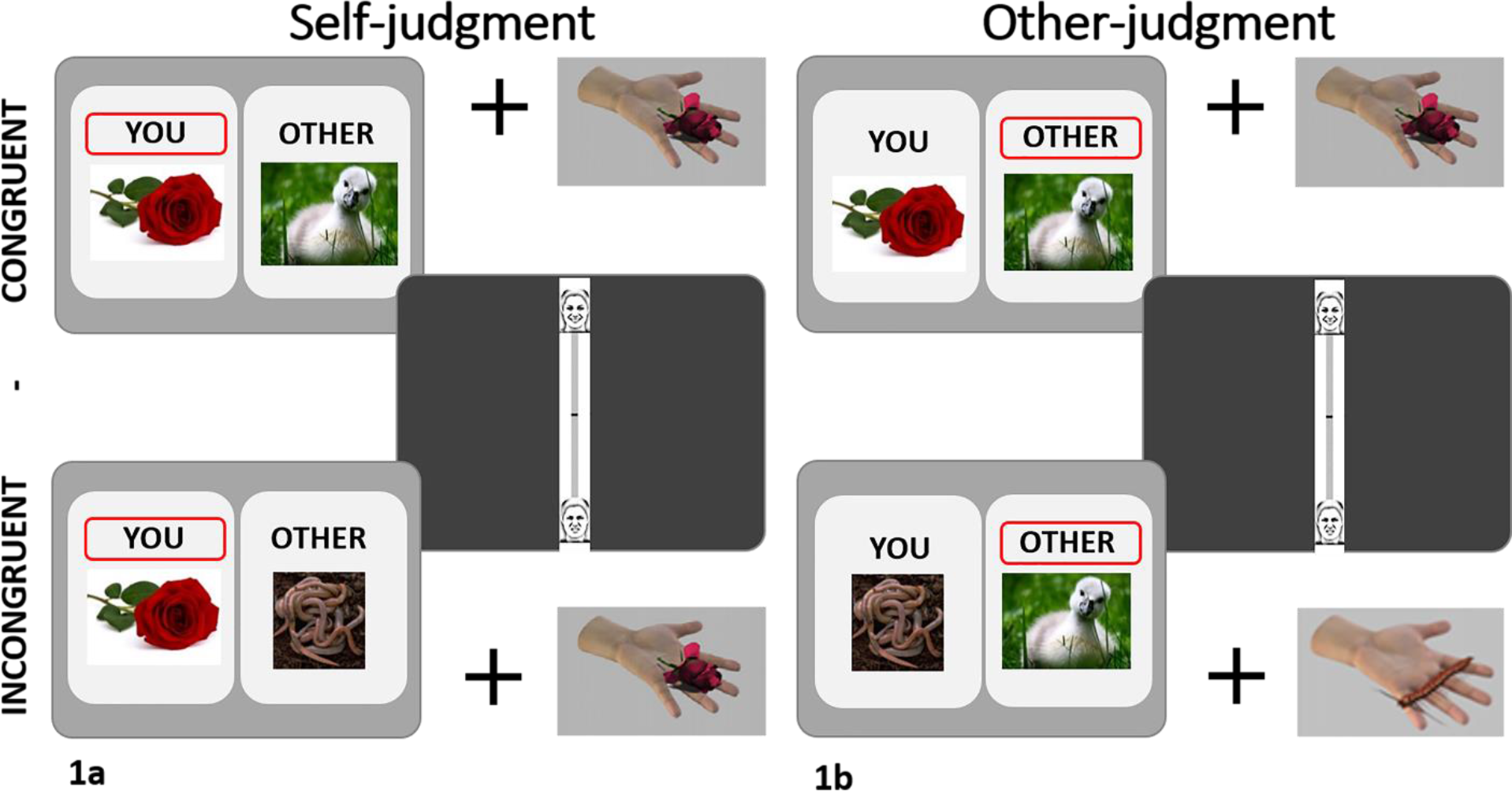
Experimental paradigm. **1a**. Self-judgment condition: participants are stroked on the palm with an object and simultaneously see on the screen an object corresponding to the touch, as well as an object indicating the touch experienced by the other participant (in reality, a confederate). The affective responses elicited in the two participants could be either congruent (upper panel) or incongruent (bottom panel). In the self-condition, participants were asked to report their own affective state during the visuo-tactile stimulation, using a visual analogue scale. **1b**. Other-judgment condition: stimulation and conditions are similar to the self-judgment condition, but participants are instructed to empathize with the other participant, and to provide ratings on their presumed affective responses.

### Computation of the EEB score

As in Silani et al. (2013) and Riva et al. (2016), EEB was computed by subtracting the ratings of the other-judgment *congruent* condition from the ratings of the other-judgment *incongruent* condition (Δ_other-judgement_), from which the corresponding difference between incongruent and congruent ratings in the self-judgment condition (Δ_self-judgement_) were subtracted. Hence, EEB = Δ_other-judgement_ - Δ_self-judgement_. Note that as in previous work, data from the unpleasant trials were multiplied by −1, so that values across the pleasant and unpleasant trials could be directly compared.

### 2.3 Additional measures

Self-report measures of depression, trait intersubjective reactivity, alexithymia, and social network were collected outside the scanner by means of questionnaires. Their purpose was to characterize better the three groups on different psychological and social dimensions and to explore the role of several possible confounding factors. Depression was measured by means of the German version (Hautzinger et al., 2009) of the Beck Depression Inventory (BDI) (Beck et al., 1961), interpersonal reactivity (personal distress, empathic concern, perspective taking, and fantasy) by means of the German version of the Interpersonal Reactivity Index (IRI) (Paulus, 2014), and alexithymia by means of the Bermond Vorst Alexithymia Questionnaire (BVAQ) (Form B; Vorst & Bermond, 2001). For the social network, three questions were administered: (1) number of friends; (2) number of close relatives; and (3) frequency of social contacts. See Table 1 for details on analyses and results.

**Table 1.** The results from the self-reported measures are presented. One-way ANOVAs were computed for the BDI (Hautzinger et al., 2009) and for each of the social network questions. In case of significance, Bonferroni-corrected pairwise comparisons were calculated to compare groups. Two multivariate ANOVAs were computed for the IRI (Paulus, 2014) and the BVAQ (Vorst and Bermond, 2001) including scales as a within-group factor (4 levels for IRI, 5 levels for BVAQ) and group as a between-group factor (3 levels), to correct for multiple comparisons of the sub-scales of each questionnaire. In case the interaction *questionnaire * group* was significant, Bonferroni-corrected post-hoc pairwise comparisons were computed for each subscale comparing the groups. Analyses were computed using SPSS v.25 (Statistical Package for the Social Sciences, IBM SPSS Inc., Chicago, IL, USA). *p-value <.05, Bonferroni post-doc test p<.05.

| Self-reported measures | Adolescents<br>Mean (SD) | Young adults<br>Mean (SD) | Older adults<br>Mean (SD) | Results |
| --- | --- | --- | --- | --- |
| BDI | 8.82 (5.63) | 8.83 (6.98) | 7 (5.004) | = |
| IRI |  |  |  |  |
| Empathic concern | 14.75 (2.27) | 16.17 (5.48) | 15.77 (2.67) | YA>AD* |
| Personal distress | 11.14 (3.44) | 15.17 (3.09) | 13.42 (2.28) | = |
| Fantasy scale | 16.07 (2.61) | 12.00 (2.49) | 11.04 (2.85) | OA<YA,AD* |
| Perspective taking | 12.79 (2.91) | 14.90 (2.44) | 15.42 (3.26) | AD<YA,OA* |
| BVAQ |  |  |  |  |
| Emotionalizing scale | 8.00 (2.67) | 7.80 (2.76) | 8.07 (2.68) | = |
| Verbalizing scale | 11.48 (3.50) | 9.40 (3.39) | 8.89 (2.94) | AD>OA* |
| Fantasizing scale | 9.50 (3.64) | 10.00 (3.40) | 12.46 (4.24) | OA > YA,AD* |
| Identifying scale | 9.54 (3.11) | 9.43 (3.12) | 6.73 (2.41) | YA,AD>OA* |
| Analyzing scale | 7.96 (2.30) | 7.77 (2.10) | 7.86 (2.55) | = |
| Social network size |  |  |  |  |
| N. of friends | 6.35 (2.70) | 6.8 (3.67) | 7.43 (4.85) | = |
| N. of close relatives | 6.55 (5.48) | 6.1 (4.44) | 7.50 (4.84) | = |
| Frequency of social contacts | 2.32 (0.75) | 2.30 (0.95) | 2.80 (0.66) | = |

### 2.4 Behavioral Analysis

An EEB score was computed for each subject and a one-way ANOVA with Group (3 levels: AD, YA, OA) as a between-group factor was then performed. In case of significant main effects, post hoc comparisons (Bonferroni adjusted) were computed. In addition, to test our hypothesis of higher EEB in adolescents and older adults, both quadratic and linear relationships between age and EEB were tested for significance. For both models the relative AIC (Akaike’s information criteria) index was computed using http://graphpad.com/quickcalcs/AIC1.cfm, which was also used to compare the different models.

### 2.5 Functional and structural MRI data acquisition, preprocessing, and analysis

Functional MRI scans were acquired using a 3T Siemens Magnetom Trio scanner equipped with a 32-channel head coil. For all participants, a high-resolution structural scan (sagittal T1-weighted MPRAGE sequence: TR: 2300 ms; TE: 2.91 ms; voxel size: 1 mm × 1 mm × 1.2 mm; slice thickness: 1.20 mm; FOV: 356 mm × 356 mm; 192 slices; flip angle: 9°), and field maps were obtained. Functional images were acquired in interleaved manner using a T2*-weighted echoplanar imaging (EPI) sequence with 33 transverse slices covering the whole brain with the following parameters: slice thickness = 3.0 mm; interslice gap = 0.3 mm; repetition time (TR) = 2060 ms, echo time (TE) = 30 ms; flip angle = 70°, field of view = 192 × 192 mm^2^; matrix size = 64 × 64. Functional MRI data were preprocessed using SPM12 (Statistical Parametric Mapping, http://www.fil.ion.ucl.ac.uk/spm). Data pre-processing included realignment and un-warping for movement artefacts, correction for geometric distortions using the acquired fieldmap, slice-time correction, co-registration of the EPI scans to the skull-stripped T1-weighted structural scan, normalization to the standard stereotaxic anatomical Montreal Neurological Institute (MNI) space, smoothing with a 6 mm full-width at half-maximum (FWHM) Gaussian kernel, and resampling of voxel size to 3 mm isotropic. The threshold used for excluding participants due to excessive motion in the scanner was fixed at 2 mm for translation and at 2° for rotation.

#### 2.5.1 Functional MRI analysis

Following the preprocessing, first-level analysis of the data of each participant was performed based on the General Linear Model framework as implemented in SPM12 (Friston, Frith, Turner, & Frackowiak, 1995). In the first-level model, eight regressors of interest convolved with SPM’s canonical hemodynamic response function were included (one for each condition of the design, i.e., pleasant incongruent self-judgment, pleasant congruent self-judgment, unpleasant incongruent self-judgment, unpleasant congruent self-judgment, pleasant incongruent other-judgment, pleasant congruent other-judgment, unpleasant incongruent other-judgment, and unpleasant congruent other-judgment), along with the corresponding eight regressors of no interest modeling the rating phase. To account for residual motion artefacts, twelve six nuisance regressors representing the realignment parameters were incorporated for each run in the first-level model as well. In line with our previous approach, which did not yield differences related to valence, both behavioral and neural data were collapsed across the two valence domains. Thus, following model estimation, the contrast of interest modelling the EEB was computed for each participant: [(pleasant incongruent - pleasant congruent) + (unpleasant incongruent - unpleasant congruent)]_other-judgment_ > [(pleasant incongruent - pleasant congruent) + (unpleasant incongruent - unpleasant congruent)]_self-judgment_, and the resulting first-level contrast images were entered in the corresponding group-level (second level) analysis. For both the functional segregation analyses and the functional connectivity we then followed the same sequential analysis approach, which consisted of three basic steps. *First*, we were interested to test, separately for each group, whether there was significant activity within the rSMG associated to the EEB contrast and whether there was rSMG connectivity with rS1, rS2 and visual cortex (VC), i.e. the areas which had shown significant activity and increased connectivity, respectively, in the young adult sample of Silani et al. (2013) (in the second, confirmatory experiment). To this purpose, we generated four masks representing rSMG, S1, S2, and visual cortex starting from the significant clusters found in Silani et al. (2013). These masks were employed for the first and the second analysis step. In this first step, we adopted a small volume correction (SVC) approach, which confined the number of statistical tests to an independently determined area for which we had strong *a priori* assumptions, thus increasing the sensitivity of the analyses. Note that this first analysis, when performed on the YA, also allowed us to assess whether we can replicate our previous findings, which had been identified in a group of similarly aged young adults. All SVC analyses used a family-wise error correction threshold of p<0.05, at voxel-level. With the *second step*, our main interest was to test differences among the three groups, both with respect to segregation and effective connectivity. Moreover, we were interested in exploring the relationship between individual differences in neural responses, and EEB. We extracted the parameter estimates for each subject for rSMG (segregation analysis) and for rS1, rS2 and visual cortex (effective connectivity), and with these values computed group comparisons, correlations with age, correlations with EEB and, in specific cases, mediation analyses. Correction for number of ROIs was not applied considered that we tested three *a priori and distinct* hypotheses, one for each area. *Finally*, while the SVC and ROI analyses of steps 1 and 2 tested activity/connectivity with higher sensitivity within predefined areas for which we had specific hypotheses, they are agnostic to potentially relevant activation/connectivity in other parts of the brain. Therefore, we complemented them with whole-brain analyses, thresholded at p<.05 FWE-corrected at voxel-level. After this description of our general analysis approach, the following paragraphs describe the specifics and the implementation of the analyses in some more detail.

##### Task-related functional segregation analyses

To test rSMG activity related to EEB we performed mass-univariate second-level random effects analyses and assessed it by means of SVC within the rSMG ROI, separately for each age group. In the next step, we tested the hypothesis of lower activity in the AD and OA groups by computing independent T-tests on the mean activity extracted from the rSMG ROI. In addition, as in the behavioral analysis, both a linear and a quadratic relation between age and EEB-related rSMG activity were assessed. Activity in rSMG related to EEB was also correlated with EEB scores. Lastly, we complemented ROIs analyses with whole-brain analysis thresholded at p<.05 FWE-corrected at voxel-level. For the whole-brain analysis we compared adolescents and older adult to young adults, in both directions (YA_EEB_ > AD_EEB_; YA_EEB_ < AD_EEB_; YA_EEB_ > OA_EEB_; YA_EEB_ < OA_EEB_).

##### Task-related effective connectivity analysis

In order to assess how rSMG connectivity with other areas of the brain differs between the three age groups, we performed psychophysiological interaction analyses (PPI, Friston et al., 1997). Following the same procedure as in Silani et al. (2013), we first extracted the deconvolved time course from the seed region rSMG (using the same mask that was used for the univariate analysis). In the second step, a PPI regressor was obtained as product of the estimated (deconvolved) BOLD signal of the seed region and the vector representing the psychological variable of interest, namely the difference between incongruent and congruent conditions in the other-judgment run. In the third step, single-subject analysis was performed by computing a GLM with the estimated neuronal activity of the rSMG, the experimental contrast and the PPI regressor. Contrast images for the PPI regressor were estimated for each subject. The analyses then followed the three steps described above. We adopted a SVC approach using the masks of the rS1, rS2 and VC to test whether we could replicate the results found in the YAs by Silani et al. (2013) and to test these same areas in the other two groups. We then conducted a ROI analysis by extracting the parameter estimates for each subject for all the three regions and then performed independent T-tests comparing AD and OA to YA as the reference group. Correlations with age were also computed. Moreover, also in this case, to explore the relation between differences in EEB and the rSMG connectivity we ran three correlations with the EEB scores. The results of this analysis suggested mediation analyses (see results). Mediation analyses were conducted using non-parametric bootstrapping procedures implemented using an SPSS Macro (Preacher and Hayes, 2008), with 5000 bootstrap resamples. Statistical significance at p < 0.05 is indicated by the 95% confidence intervals not crossing zero. As a last step, whole-brain analysis was performed and group comparisons were computed, again comparing AD and OA to YA in both direction.

#### 2.5.2 Structural MRI analysis

In addition to functional segregation and connectivity analyses, we analyzed the structural MRI data by means of voxel-based morphometry (VBM) analyses (Ashburner and Friston, 2000), in order to investigate age-related differences of rSMG and other areas. Analysis of gray matter volume was performed via voxel-based morphometry implemented in the CAT12 toolbox (http://dbm.neuro.uni-jena.de/cat/) within SPM12. Preprocessing included bias field correction, segmentation in gray matter, white matter and cerebrospinal fluid using a segmentation approach based on adaptive maximum a posterior segmentation and partial volume segmentation. The resulting segmentations were normalized into Montreal Neurological Institute (MNI) space using Diffeomorphic Anatomic Registration Through Exponentiated Lie algebra algorithm (DARTEL; Ashburner, 2007) with the DARTEL MNI template image included with the CAT12 toolbox. The segmented, normalized and modulated images reflecting gray matter volume were finally smoothed with an 8 mm FWHM Gaussian kernel and used for subsequent ROI statistical analysis. ROI analysis was then performed using the rSMG mask also used for the functional analyses. Group comparisons were computed to explore differences between AD and YA and between OA and YA. A correlation analysis of the relationship between age and rSMG volume was also performed. Total intracranial volume (tiv) was included as covariate of no interest in the models. As a last step, group comparisons at whole-brain level were computed also for the structural data.

## 3. Results

### 3.1 Behavioral results

The one-way ANOVA computed on EEB scores revealed a main effect of group (F (2,91) = 21,395; p<.001; partial *ɳ*^*2*^ = .325). Post-hoc comparisons revealed that OA had a significantly higher EEB than both YA and AD (all p<.001) (AD_mean_ = .256, AD_SD_ = 1.068; YA_mean_ = .047, YA_SD_ = 1.622; OA_mean_ = 4.834, OA_SD_ = 5.618), see Fig. 2a. Contrary to our hypothesis, though, no differences between YA and AD were found. Both the tests of the linear and a quadratic relation between EEB and age were significant (Linear: F(1,91) = 49.970, p < .001, R^2^=.357; AIC = 167.58; Quadratic: F(2,91) = 28.914, p < .001, AIC = 178.75). However, the comparison of the two models by means of the AIC index indicated that the linear model is 266.91 times more likely to be correct than the quadratic model (Fig.2b).

**Fig. 2.**
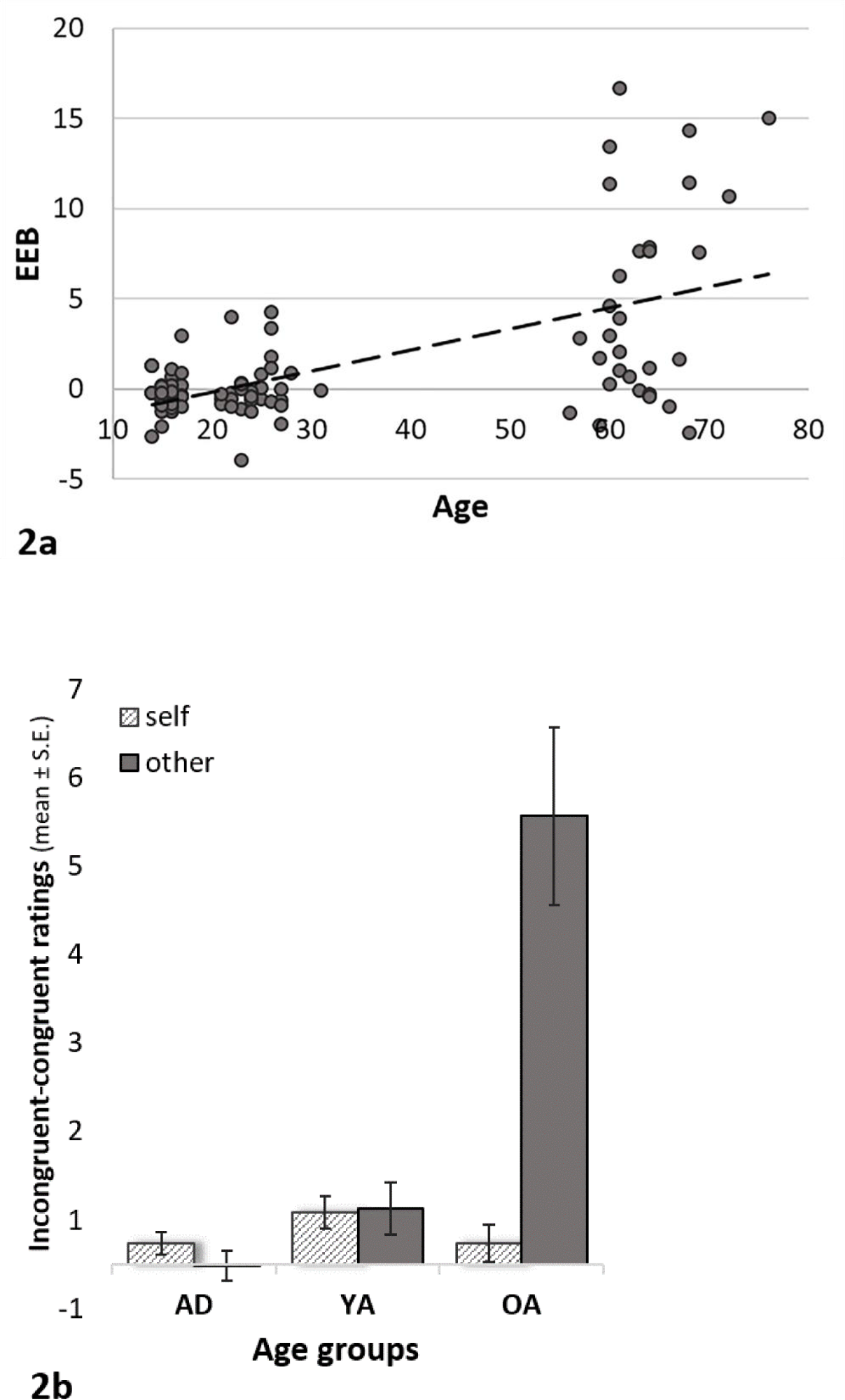
EEB at different ages. **2a**. Scatterplot of individual EEB (y-axis) values in adolescents, young adults and older adults (age on x-axis). **2b**. Differences between ratings provided in incongruent vs. congruent trials, separately for self- and other-judgment conditions. While no group differences emerged in the self-judgment condition, older adults showed, compared to the other two groups, a significantly higher incongruity effect in the other-judgment condition, giving rise to a higher EEB (see text).

**Fig. 3.**
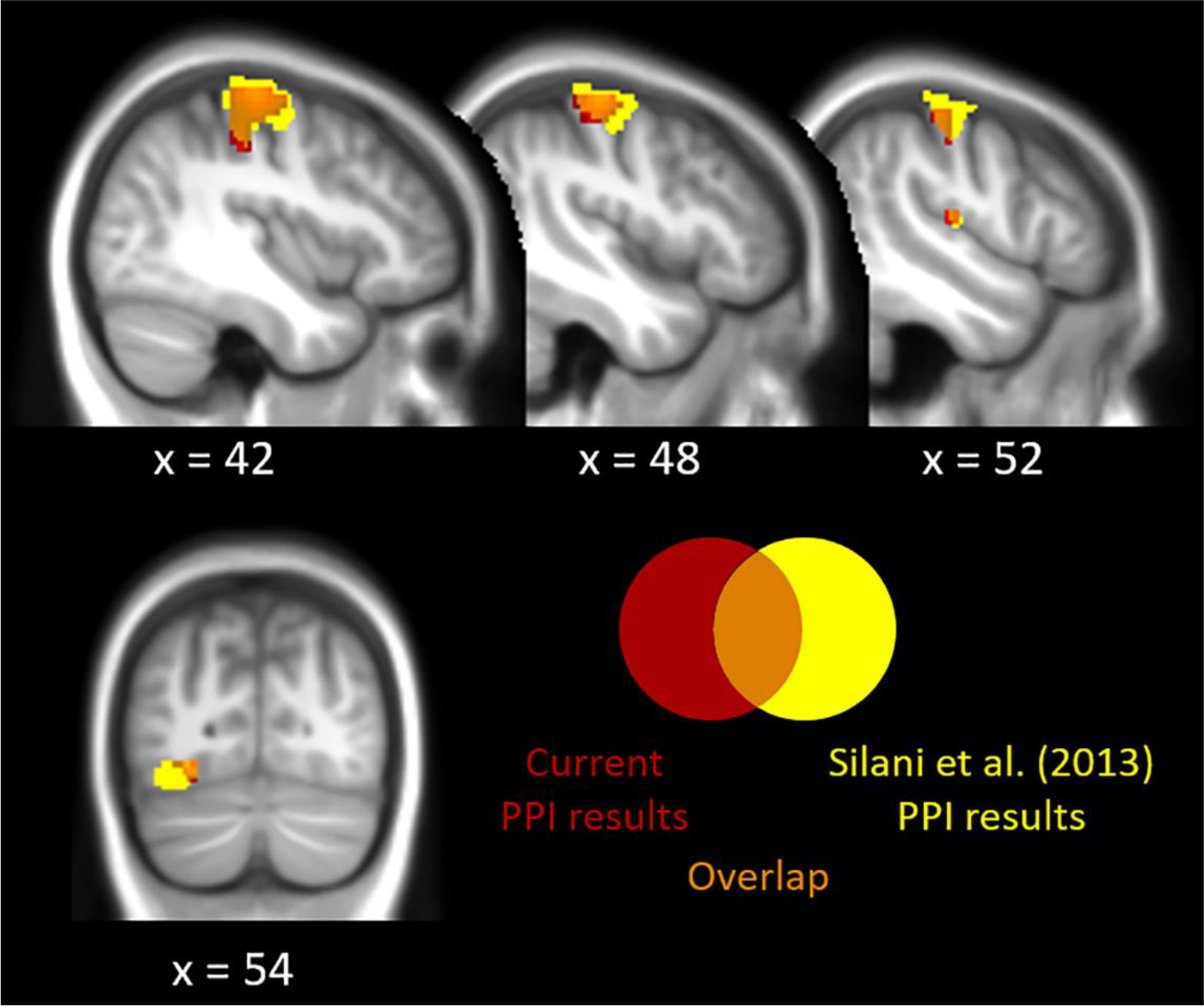
Brain regions showing increased effective connectivity (PPI analysis) with the rSMG in young adults: S1, S2 (sagittal view) and VC (coronal view) (p=.005 FWE). These results (in red) replicated previous results (in yellow) (Silani et al. 2013), as indicated by the orange overlap between the present and the previous study.

**Fig. 4.**
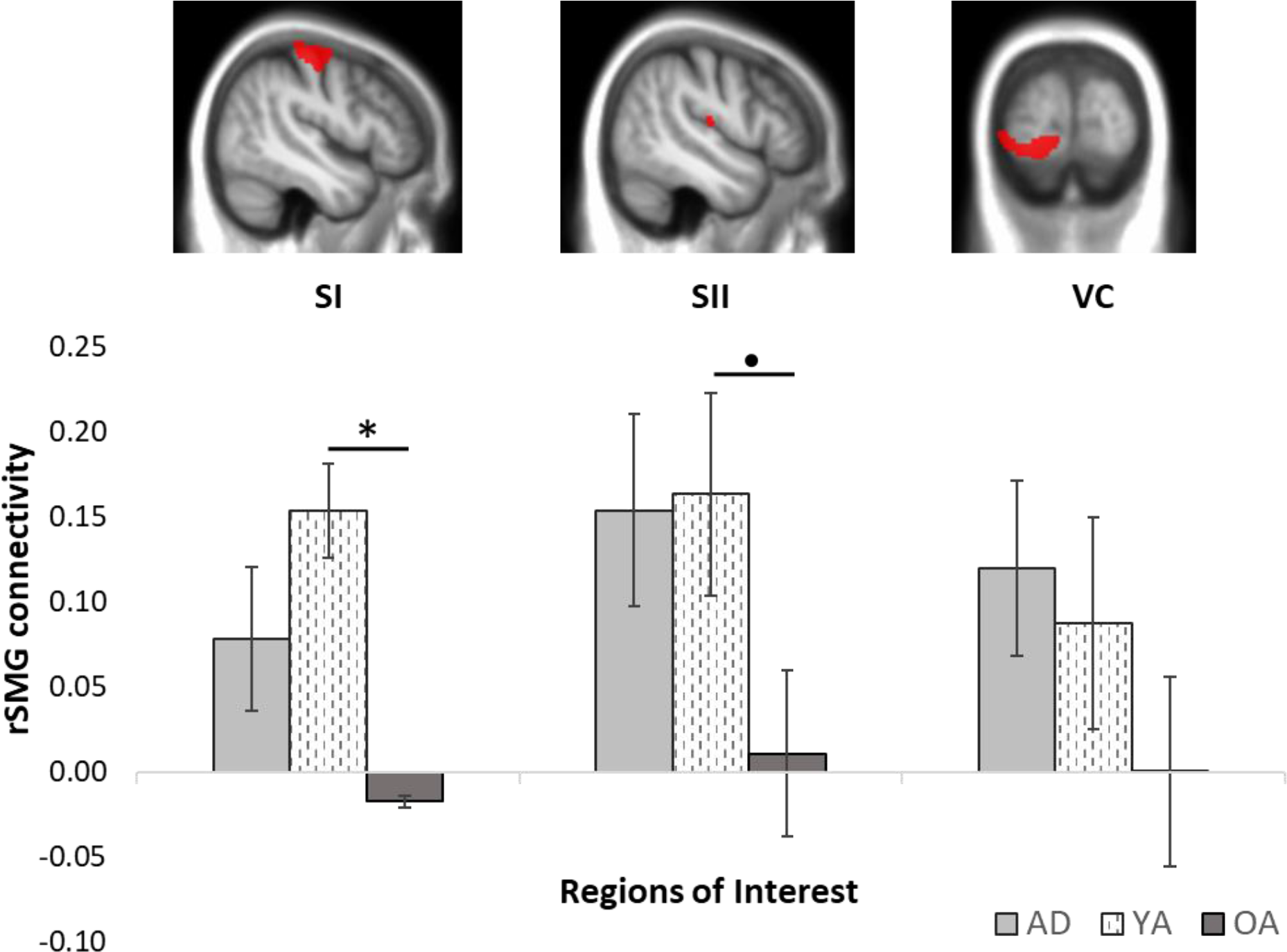
Group differences of effective connectivity between rSMG and S1, S2 and visual cortex (VC). Older adults showed lower effective connectivity of the rSMG with S1 and S2 compared to the young adults, while no differences emerged between adolescents and young adults. The top of the figure shows the location and the extent of the ROI masks, built from the corresponding results in Silani et al. (2013) (see text for details).

### 3.2 Functional MRI results

#### 3.2.1 Task-related segregation results

The SVC analyses within rSMG showed significant activity in the rSMG in YA (peak voxel at MNI x/y/z= 60/−34/40), and thus replicated the findings of Silani et al. (2013). No significant voxels were found, however, in the AD and in the OA group. Testing differences in rSMG activity among the three groups revealed no group differences, nor was there any significant correlation, neither linear nor quadratic, between EEB-related rSMG activity and age. The complementary whole-brain analyses revealed significant activity in the anterior mid-cingulate cortex (aMCC, −3/−4/46) in YA, but no group differences in this area either. However, group comparisons revealed greater activation of the right temporal pole (36/11/−23) in the OA compared to YA.

#### 3.2.2 Task-related effective connectivity results

In the YA group, significantly increased connectivity between rSMG and rS1 (45/−22/64), rS2 (48/−19/16) and visual cortex (V4 according to anatomy toolbox,−27/−76/−8) were largely in line with the results of Silani et al. (2013). In the same vein, AD showed significantly increased connectivity between rSMG and rS1 (42/−7/55), rS2 (51/−19/16) and the visual cortex (medial/left hemisphere, −15/−91/−17). On the contrary, OA did not show any significant changes in rSMG connectivity. Comparisons across groups revealed that OA, compared to YA, showed significantly lower connectivity of the rSGM with rS1 (t(59) = 2.227, p = .030), and a trend towards lower connectivity of the rSMG with rS2 (t(59)= 1.936, p =.058). No differences were found between AD and YA. Moreover, a negative correlation was found between age and connectivity of the rSMG with rS1 and rS2 (S1: r=−.222 p=.034, S2: r=−.303, p=.003). In order to go deeper in the relation between age, EEB and rSMG connectivity, we performed two mediation analyses with age as the continuous predictor, EEB as the outcome variable, and in the first one the connectivity between rSMG and rS1 as the mediator, and in the second one the connectivity between rSMG and rS2 as the mediator. The analyes revealed that the connectivity between rSMG and rS1 did not mediate the effect of age on the EEB, while we found a significant mediation effect of the connectivity between rSMG and rS2 with respect to the relation between age and EEB (indirect effect=.008, SE=.0038, 95% CI=[.0012, .0158]). However, the mediation was only partial since the direct effect of age on EEB was significant as well (p<.05). Lastly, whole-brain analyses revealed a decreased connectivity associated to increasing age for right visual cortex (21/−88/28), aMCC (−3/26/ 40), and midbrain (3/−16/−2).

### 3.3 Structural MRI results

Group comparisons revealed that AD have greater gray matter volume than YA (t(60) = 3.513, p = 001) and YA greater than OA (t(59) = 3.513, p <.001. A significant negative correlation also emerged between age and rSMG gray matter volume (r=−.408, p<.001). Whole-brain analyses, controlling for total intracranial volume, showed, at an FWE-corrected voxel-level threshold of p<.05, higher gray matter volume for AD than YA in a number of clusters, among others and most prominently in the superior medial frontal gyrus, the right angular gyrus/temporoparietal cortex, the medial parietal cortex including the precuneus and the posterior cingulate cortex. YA had more gray matter than OA in all of these regions and in additional extended clusters, including medial occipital cortex, bilateral cerebellar regions, lateral frontal, parietal and temporal regions, the bilateral insula and subcortical regions. No significantly higher gray matter volume were found in OA compared to YA and in YA compared to AD.

## 4. Discussion

We used a multi-level neuroimaging approach, combining morphometry, functional segregation and task-related connectivity analyses, to shed light on the neural underpinnings of age-related differences in emotional egocentricity. Our main findings include a) the replication of the involvement of rSMG and its connectivity in overcoming emotional egocentricity in young adults; b) the partial replication of previous behavioral findings, that adults over around 60 years show higher EEB than young adults, from whom adolescents (contrasting our prediction) do not differ; c) the association between age-related changes in EEB and changes in rSMG effective connectivity with somatosensory cortices, subtended by structural changes in rSMG across the lifespan.

More in details, the previous finding (Riva et al., 2016) that older adults are more emotionally egocentric than young adults was replicated, whereas adolescents did not show higher EEB than the young adults – thus being in contrast with our predictions.

With regard to the neural underpinnings of emotional egocentrism in young adults, results from both the functional segregation and the effective connectivity analyses replicated previous findings reported in a sample of similar age (Silani et al., 2013): Young adults showed indeed significant activity in the rSMG related to EEB, and increased effective connectivity between rSMG and rS1, rS2, and the visual cortex. This confirms the view that rSMG is an area crucial for reducing EEB, and that this is achieved by connecting with sensory-perceptual areas to implement self-other distinction (see also Kanske et al., 2015; Hoffmann et al., 2016). The complementary whole-brain analysis revealed that overcoming EEB was also related to significant activity in the aMCC, an area (amongst others) involved in task monitoring, conflict resolution (Weissman et al., 2003; Kim et al., 2011), and affect regulation in the domain of empathy (Lamm et al., 2019, for review).

With regard to the older adults, no significant EEB-related rSMG activity emerged, though direct comparison between young and older adults did not reveal a significant difference. However, significant lower connectivity between rSMG and rS1 and a trend towards significant connectivity with rS2 was observed, compared to young adults. Mediation analysis revealed the connectivity of rSMG with rS2 to be a partial mediator of the relationship between age and EEB, uncovering the relevance of the task-related rSMG connectivity in overcoming EEB. These findings might be related, at the structural level, to the observation of smaller gray matter volume in the rSMG compared to the young adults. Whole-brain analysis moreover showed that older adults, compared to young adults, presented significantly higher activity in the temporal pole, an area involved, among other things, in socio-cognitive processes (Pehrs et al., 2017). A compensatory mechanism of the older brain dealing with less efficient brain activity (Cabeza et al., 2002; Reuter-Lorenz and Cappell, 2008) might be at the origin of this result. Note though that this result was not predicted *a priori* and should be regarded as exploratory.

On the other end of the lifespan, adolescents did not display themselves as more egocentric than young adults. Different reasons may account for the deviation from the present from our previous findings. First, in the previous (behavioral) study ratings had been collected using a response device (a touch screen) that enabled faster and more automatic responses. In the present MRI study, responses had to be collected by moving a cursor on the response scale. The possibility to adjust the cursor position while entering the response might resulted in additional time and reflection to overcome and control for initial bias. Note that this discrepancy between response modalities has also been reported in our previous studies (Silani et al., 2013) in young adults, and also may account for a lack of significant EEB in the YA group. Another reason might be a difference in the two adolescent samples, which, amongst general sampling issues, might be due to possible cultural and educational differences, as data were collected in different countries (Italy and Austria). At the brain level, adolescents showed greater gray matter volume in rSMG compared to young adults, which is likely to indicate ongoing development processes, such as pruning and myelination, as proposed by various researchers (Ducharme et al., 2015; Tamnes et al., 2017). In line with such ongoing development and differentiation of the rSMG, no significant activity in this area was found for the EEB contrast; although, as for the older adults, no significant differences emerged between adolescents and young adults. Notably, and differently from the older adults, no significant differences occurred between adolescents and young adults in rSMG effective connectivity. Indeed, significant task-related connectivity between rSMG and rS1, rS2 and visual cortex was also observed in the adolescents.

Considering data from both adolescents and older adults, age-related changes in rSMG connectivity seems to play a central role in changes of emotional egocentricity across different ages. When considering the whole sample, increasing age has been found to negatively correlate with connectivity between rSMG and somatosensory cortices, and in particular the coupling between rSMG and rS2 partially mediates the relationship between age and EEB. S2 is a brain area involved in a variety of processes, from the perception of touch intensity (Case et al., 2017) to emotional processing (Adolphs et al., 2000) and attentional modulation of somatosensory stimuli (Chen et al., 2008). Importantly, in addition to being activated by first-person touch stimulation, S2 has been associated to observation of vicarious touch (Keysers et al., 2004) and to empathy for touch (Jackson et al., 2006). Thus, a possible, though speculative, interpretation might be that both self- and other-related emotional experiences are represented in S2 and transferred to rSMG. In young adults, when the two representations are incongruent, the coupling between S2 and rSMG increases, possibly because of the higher complexity/greater quantity of information exchanged with rSMG.

Being a central area for self-other distinction in the emotional domain (Silani et al., 2013; Steinbeis et al., 2014), rSMG keeps separated and weights information related to one’s own and to the other’s emotional states, providing the basis of the empathic judgment. However, since the increase in the rSMG-rS2 coupling is not observed in older adults, this might suggest that the complexity associated to simultaneous incongruent emotional states between self and other is not transferred to the rSMG. Thus, participants in this group may use the more salient representation (i.e., the self) to inform their empathic judgment resulting in a higher egocentric bias.

The relation between age and task-free functional connectivity (e.g.: resting state) have been fairly extensively investigated (Geerligs et al., 2014, 2015; for a review on aging: Damoiseaux, 2017; McCormick et al., 2018) and showed associations between age-related differences in social/cognitive abilities and age-related changes in task-free functional networks (e.g.: default mode network). However, less investigated is the relation between age-related changes in socio-cognitive processes and differences occurring with development and aging in task-related effective connectivity. The current study provides an example of how differences in regional functional activity might not always be able to account for age-related differences in (socio)cognitive processes, whereas task-related functional connectivity might play a key role in identifying these differences. Thus, a more systematical and regular analysis of task-related effective connectivity in investigating age-related differences in brain functionality seems advisable for future investigations. Despite several strengths, there are also some limitations that need specific consideration. First, the confederate playing the “other” in the task was a young adult for all the three groups and this might have influenced the degree to which the participants were able to empathize with them. In this respect it is important to note that the present sample (as reported in Riva et al., 2018) did not reveal any behavioral indications of increased difficulty to empathize, although we did observe differences in empathy-related anterior insula activity. Future studies are thus needed to test whether the present findings generalize to empathy with age-matched persons. Second, as in any cross-sectional study, factors associated to cohorts rather than age might have played a role. Third, despite our paradigm has been already employed in different studies/experiments (Silani et al., 2013; Tomova et al., 2014; Riva et al., 2016), the extension of the present results to other types of paradigm investigating the EEB (Steinbeis et al., 2014; von Mohr et al., 2019) is required. Finally, the use of a female sample, for the reasons outlined above, requires further research to test whether the present results extend to the male population.

## Conclusions

The current study confirms that older age is associated with higher EEB than in young adults. Whether or not adolescents show higher EEB remains controversial considering on the non-replication of previous findings. We also corroborated that, in young adults, rSMG is a central area for efficient self-other distinction in the emotional domain. At the same time, age-related differences in emotional egocentricity seem to be better explained by differences in the effective connectivity of the rSMG with somatosensory cortices, and especially with rS2. Taken together, the present and previous findings suggest that rSMG works in interaction with other, predominately sensory-perceptual brain areas to integrate as well as to differentiate affective information pertaining to self and other.

## Acknowledgments

The study was supported by the Austrian Science Fund (FWF, P 29150 to CL).

